# Emotional reactivity to aversive primes impedes motor preparatory activity in functional neurological disorders

**DOI:** 10.64898/2026.04.16.718849

**Authors:** Viridiana Mazzola

## Abstract

Patients with functional neurological disorders (FNDs) show impaired control of voluntary actions in the absence of organic neurological damage. The inconsistency between objective neurological clinical signs and actual performance of the same movements in slightly different contexts points to an abnormal self-focused attentional role towards movement execution. Yet, it remains unexplained what triggers a higher level of self-focused attention in FNDs and how this interferes with voluntary movements. Given the known threat sensitivity manifested by patients with FNDs, I hypothesized that under negative affective conditions self-focused attention might be heightened in FNDs in an automatic way so as to impede the execution of a voluntary action. Specifically, I used fMRI to investigate effective brain connectivity in “self-referential” and “limbic” circuits to delineate the causal functional architecture accounting for the FND specific activity when preparing a movement under aversive conditions with different levels of emotion awareness. Seventeen FND participants and seventeen healthy volunteers performed a motor task (key press and release) after having been exposed to an aversive or neutral picture prime using a sandwich mask paradigm. Behaviorally, the FND group had showed a slower reaction time across all task conditions and a high rate of missing key-press responses following associated to aversive primes. Dynamic Causal Modeling (DCM) analyses showed that the FND group emotional information did not engage a limbic network as observed in the healthy control group, but rather a different self-referential associated network. In this functional architecture, the aversive masked condition exerted a direct inhibitory effect on forward connections between the left IFG and left precentral motor cortex. These findings show how affective processing can impact on voluntary motor control in FND, helping to reduce the explanatory gap between emotionality and readiness to act as a potential process of functional motor symptom production.

## Introduction

People with functional neurological disorders (FNDs) present impaired voluntary motor control despite a lack of neurological disease and of a clear structural brain basis affecting their sensorimotor system (Espay et al. 2018, Bègue et al. 2019). In everyday life contexts, often in an unpredictable way, they appear unable to control simple voluntary actions, like walking or moving an arm or a leg, or stopping a tremor or a dystonic posture, manifestations that can also fluctuate over time. In addition, they report a frustrating and frightening experience of feeling out of control of their own movements, complaining also for a high level of variability of the triggering conditions, intensity, and duration.

The well-known inconsistencies between neurological clinical signs (muscle power or movement speed) and performance of the same movements in slightly different contexts let the neurologists suppose an underlying attentional role towards the impaired movement. Actually, it can occur during their neurological examination that if the patient’s attention is directed away from the abnormal movement, there is almost a normalization of that movement (Kennedy et al. 2007). In contrast, the reverse pattern is manifested in patients with organic neurological signs. Experimental investigations also support the role played by attention and expectations in FNDs (Roelofs et al. 2003, Pareés et al. 2012, 2014a). The FNDs model of Edward and colleagues seem to go in this direction as well (Edwards et al. 2012). Within a predictive coding approach, they hypothesize that the FNDs arise when the precision of abnormal intermediate-level expectations is enhanced by a misdirected, self-focused attention. As a flexible approach to FND causation, it could account for the variability of the functional motor (and sensory) symptoms. Nevertheless, it still remains unexplained what unexpectedly triggers abnormal self-focused attention in people with FNDs and how this interferes with their voluntary movements, leaving also the clinicians with very limited scope of intervention. On one hand, a different approach to the neural motor functioning shifts the focus from the meaning of the motor output to the nature of the dynamical system that creates the required, precisely patterned, command (Shenoy et al. 2013). According to this dynamical systems view, the preparatory activity serves not to tune to movement parameters (e.g., reach endpoint or velocity), but rather to initialize a dynamical system whose future evolution will produce movement (Churchland et al., 2010). Indeed, the preparatory activity is predictive of reaction time and movement variability (Bastian et al., 2003; Churchland et al., 2006a; Churchland et al., 2006b; Riehle and Requin, 1993), and its disruption delays the movement onset (Churchland and Shenoy, 2007). On the other hand, from a psychological approach a potential underlying process might concern the link between a well-known higher threat sensitivity manifested by these patients and their readiness to act (Arciero and Bondolfi, 2009; Bakvis et al. 2009, 2010; Blakemore et al. 2016). Indeed, previous neuroimaging studies showed an increased functional coupling between the amygdala and the supplementary motor area in participants with FNDs when viewing both fearful and happy facial expressions compared with healthy volunteers (Voon et al. 2010).

All the above considerations, together with my clinical practice with FNDs as a psychotherapist,lead to the hypothesis that under certain emotional conditions, especially the threatening ones, these patients react with heightened self-referential attention, in such an automatic way that it can somehow interfere with voluntary movement control.

Therefore, in the present study I investigated whether and to what extent a different emotionality would affect the preparatory motor neural activity in the FND patients compared to healthy volunteers. A functional neuroimaging was used during a novel paradigm allowing to investigate how motor action is modulated by aversive priming, in both FND and healthy individuals. Seventeen FND participants and seventeen healthy controls underwent an fMRI session where they performed a double motor task (sustained key press and release) after having being exposed to an aversive or neutral picture prime, presented either supraliminally or subliminally using a sandwich mask procedure. Three functional neural architectures (two emotion-related limbic circuits with and without the amygdala and one self-focused circuits) were then compared to check which among them would be more likely to exert an excitatory or an inhibitory effect on motor systems in FND patients and healthy participants, while preparing and executing hand movement. Indeed, using a dynamic causal modeling analysis (DCM), I obtained statistical estimates of which connectivity model offered the optimum balance between simplicity and fit to the data using Fixed Effects Bayesian Model Selection family inference analysis (FFX BMS) (Friston et al., 2003; Penny et al. 2010), followed by a model parameter estimates analysis of Bayesian Parameters Averaging (BPA) (see Methods). I expected that self-referential attention triggered by aversive emotional primes would exert a distinctive inhibitory modulatory effect on functional connectivity within the motor action module in FND participants, but not healthy controls.

## Methods

### Participants

Seventeen right-handed FND participants and seventeen healthy controls, matched for age and gender, were enrolled. The FND participants were recruited at the Geneva University Hospital. The diagnosis was established according to DSM-5 criteria of Conversion Disorder (Functional Neurological Symptom) code F44.4-5-6-7 and a board-certified neurologist confirmed the presence of positive functional features (DSM criteria B) (Tab.1). Exclusion criteria included age <18 years old, a history of drug or alcohol abuse, previous head trauma with loss of consciousness, severe psychiatric disorder (depression with suicidality, acute psychotic symptoms) and co-morbid neurological disease, and any other significant medical condition. The study was approved by the ethics committee of the University Hospitals of Geneva. Informed written consent was obtained from all participants before their participation.

**Table 1:**
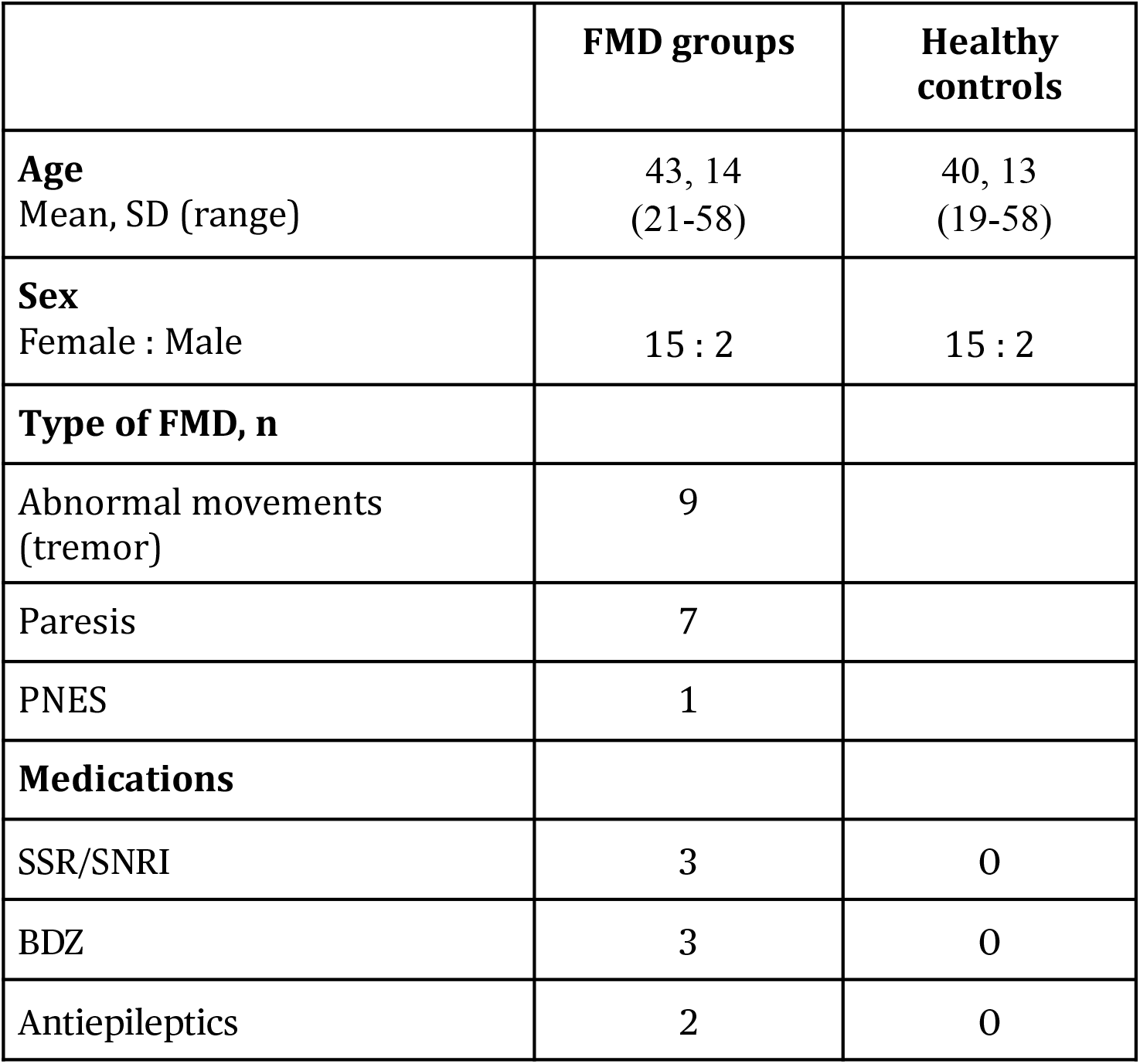
Demographic and clinical characteristics of the participants.

### Questionnaires

In order to evaluate whether the two groups would had significant differences in level of anxiety and depression, the State Anxiety Inventory (STAI-S) (Spielberger, 1983) and the Beck Depression Inventory (BDI-II) were administered.

### fMRI experimental paradigm

#### fMRI task

The functional MRI session consisted of a single run of an event-related design task I designed. The primes consisted of sixty colored pictures of different objects and scenes selected from the International Affective Picture System (IAPS) (Davis et al. 1995), with two different affective contents (aversive and neutral). They were shown with a sandwich masking technique in which masks were created from the same images using the “liquefaction” function in Adobe Photoshop (generating random swirls to degrade the image), presented in rapid succession both before and after the prime picture. The task involved four experimental conditions: either aversive or neutral primes, shown either in masked or unmasked conditions. The experiment consisted of 120 trials (30 per condition), each with a duration of about 10s. Each trial started with a fixation cross lasting for 1500 ms. Then, in the masked condition, a blank (100ms duration), three masks (100ms duration), a prime (33ms duration), and three masks (100ms duration) were presented sequentially, efficiently preventing conscious perception of the prime. In the unmasked condition, the prime lasted for 100ms instead of 33ms (Fig.1). Each prime picture was repeated twice, according to the masked or unmasked condition, in a counterbalanced and randomized order. At the end of prime and mask sequence, a picture of a landscape was displayed and remained on the screen for a variable duration (mean 6 sec, jittered between 4 sec and 8 sec). Participants were asked to look carefully at the sequence of images and to press the button as soon as the landscape picture appeared, and then maintain the pressure and release the button as soon as the landscape picture disappeared.This allowed us to probe both motor execution (key press) and motor inhibition (key release). The time window to record responses was set at 3s. Before the scanning session, participants were trained outside the scanner to perform a brief version of the task with six trials in each condition (using different prime pictures). The total scanning time was about 16 minutes. All the participants performed the task using their right hand. The stimulus presentation and response recording was controlled by Cogent 2000 (Welcome Department of Imaging Neuroscience).

**Fig. 1:**
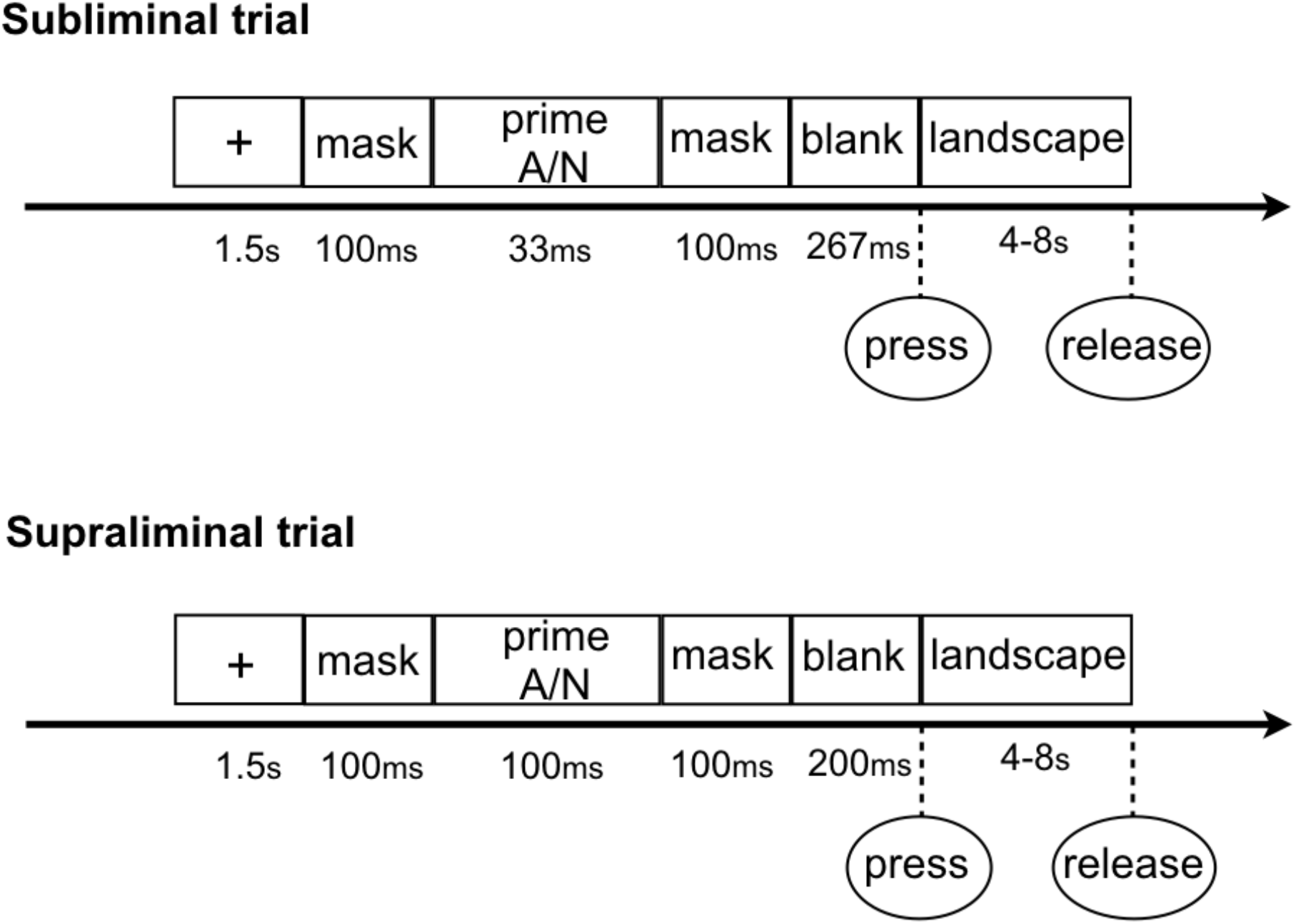
Schematic illustration of the fMRI task using a sandwich mask procedure.

#### Behavioral analyses

A repeated-measures ANOVA between groups with prime type (aversive and neutral) and conscious modality (masked and unmasked) as within factors was performed both on key press and release reaction times (RTs). Missing responses (3sec before and after event onset) were not included within the analysis of RTs. The distribution of missing responses was also computed to evaluate any differential pattern of actual motor action across conditions, and then submitted to Pearson Chi square tests to compare groups and conditions. Accuracy was computed in terms of the absolute number of misses. Questionnaires and behavioral data were analyzed with SPSS22 (Inc., 2009, Chicago, USA).

#### fMRI data acquisition and analyses

Scanning was performed at the Brain & Behaviour Laboratory (BBL), managed by the Swiss Affective Science Center at UNIGE. All functional data were acquired using a slice-accelerated multiband gradient-echo echo planar imaging (EPI) sequence using T2* weighted blood oxygenation level dependent (BOLD) contrast (multiband acceleration factor: 3, GRAPPA acceleration factor: 2). This allowed for a whole-brain coverage with 66 slices and an isotropic voxel size of 2 × 2 × 2 mm with the following parameters: repetition time (TR) = 1300 ms, echo time (TE) = 25 ms, flip angle = 60°, slice thickness = 2 mm, FOV = 192 × 224 mm. A total of 680 neutral EPI volume images were acquired. In addition, a high-resolution T1-weighted anatomical scan was performed for each participant (MPRAGE, TR=1900ms, TE=2.27ms, TI=900ms, flip angle=9°, 192 sagittal slices, voxel dimensions=1mm isotropic, 256×256 pixels, PAT factor=2).

##### Preprocessing

Data were preprocessed and analyzed using statistical parametric mapping SPM12 (Wellcome Department of Cognitive Neurology, London, UK). All images from the three runs were realigned, corrected for slice timing, normalized to the MNI space (reslicing 3×3×3mm voxel-size), and spatially smoothed with an 8-mm full width at half-maximum (FWHM) Gaussian kernel. Structural and functional images were spatially normalized into a standardized anatomical framework using the default EPI template provided in SPM12, based on the averaged-brain of the Montreal Neurological Institute and approximating the normalized probabilistic spatial reference frame of Talairach and Tournoux (1988). High-pass frequency filter (cutoff 128s) and corrections for auto-correlation between scans were applied to the time series. Individual events were modeled by a standard synthetic hemodynamic response function (HRF). A fixed-effect model at a single-subject level was performed to create images of parameter estimates, which were then entered into a second-level random-effects analysis.

##### Statistical analyses

I performed two parallel but identical statistical analyses on the functional data for the whole-brain and cerebellar normalized images. Eight event-types were defined per subject, corresponding to each condition of interest. The condition of interest were: aversive masked (Am), aversive unmasked (Au), neutral masked (Nm), neutral unmasked (Nu), landscape following Am, landscape following Au, landscape following Nm, and landscape following Nu. No-response trials were modeled as regressor of no interest. Then, eight contrast images corresponding to these conditions were created using one-sample t-tests in all subjects and then entered at the second level into a repeated-measures 2×8 ANOVA (flexible factorial design implemented in SPM12). For whole-brain analyses the statistical threshold was P=0.05 FWE corrected. In order to verify the research hypothesis I carried out interaction analysis an interaction analysis between groups and type of primes ([Am+Au]-[Nm+Nu]). To verify from which group the resulting effects were driven, I inclusively masked the resulting SPM with group specific main effects of factor. Anatomic and Brodmann areas labeling of cerebral activated clusters was performed with the SPM Anatomy Toolbox (Eickhoff et al. 2005).

##### Cerebellar normalization

I used a separate normalization process for data from the cerebellum. The registration between individuals and MNI space is suboptimal in the cerebellum when using a standard whole-brain normalization process (Diedrichsen et al., 2009). Because cerebella vary relatively little between individuals compared with the cortical landmarks used for whole-brain normalization, it is possible to achieve a much better registration by normalizing the cerebella separately. Moreover, precise spatial registration is important because cerebellar structures are small relative to cortical structures. To this aim, I used the SUIT toolbox (Diedrichsen et al., 2009) for SPM12 allowing us to normalize each individual’s structural scan to an infratentorial template, and then used the resulting deformation maps to normalize the cerebellar sections of each person’s functional images. The SUIT toolbox has the additional advantage that coordinates can be adjusted from MNI space to the corresponding coordinates on the unnormalized Colin-27 brain, which is described anatomically in a cerebellar atlas. I used this feature to identify anatomical regions within the cerebellum.

### DCM analysis

#### VOI extraction

For each region included in the DCM analysis, the individual BOLD time-series were extracted as the first eigenvariate of all significant voxels within a 6 mm radius sphere centered on local maxima in MNI coordinates, as defined from the corresponding main effect and interaction contrasts in the initial whole-brain SPM analysis [dmPFC: -6 52 20; PCC: 8 -40 14; vmPFC: -8 28 -16;amygdala: 20 -2 -12; IFG: -50 12 24; precentral gyrus: -36 12 40, 62 2 14; SMA: 0 40 36, -12 10 52].

#### DCM model space

DCM analyses were performed with the DCM12 routine implemented in SPM12. I defined the model space based on the research question: which module of emotional reactivity does better drive hand preparatory activity in FND and healthy groups? Indeed, when facing an aversive scene, the ventromedial prefrontal cortex/vmPFC and the amygdala can be expected to integrate a limbic subnetwork modulating online a fear response (Bzdok et al 2013). On the other hand, the dorsomedial prefrontal cortex/dmPFC and posterior cingulate cortex/PCC subsystem is preferentially engaged when participants make self-referential judgments about their present situation or mental states (Andrews-Hanna et al. 2010). In fact, the PCC plays a more active role in controlling the balance between an internal and external focus of attention (Leech et al, 2011), while the dmPFC is directly involved in the evaluation of externally generated social information (Murray et al. 2012). For the action hand module I selected bilateral supplemental motor areas (SMA) and bilateral precentral gyri (Grefkes et al. 2008). According to role play by the left inferior frontal gyrus/IFG during hand action planning to use tool and execution (Cha et al. 2016), I added it as the node in the action hand module. For all models, I defined fixed connections based on anatomical evidence and then compared these different architectures in both healthy controls and patients. In all cases, we defined a motor action module including motor and premotor regions in both hemisphere (ROIs in precentral gyri, bilateral SMA, and left IFG) and added modulatory regions associated with limbic emotional processes and self-focus processes (Fig. 2). We first divided our models in three families (family#1: “vmPFC”; family#2: “vmPFC-amygdala”; family#3: “dmPFC-PCC”) based on the driving input into the network. Specifically, the family#1 incorporated connections from vmPFC to the motor action module. The family#2 used the vmPFC-amygdala as modulation sources instead, whereas the family#3 replaced the vmPFC-amygdala with the dmPFC and PCC. Then I specified 12 generative models for the first family#1, 26 models for family#2, and 15 models for the family#3. In all of them I varied regions as driving input and probed for any modulation of both the regions and its connections as a function of the different prime conditions (aversive vs neutral scenes with masked vs non-masked presentations). In all models, the first region received all the conditions of the session as a driving input. The connections within the motor action module were fixed as expected for the right hand motor activity (Grefkes et al. 2008). All models were two-state nonlinear (Churchland et al. 2010).

**Fig. 2:**
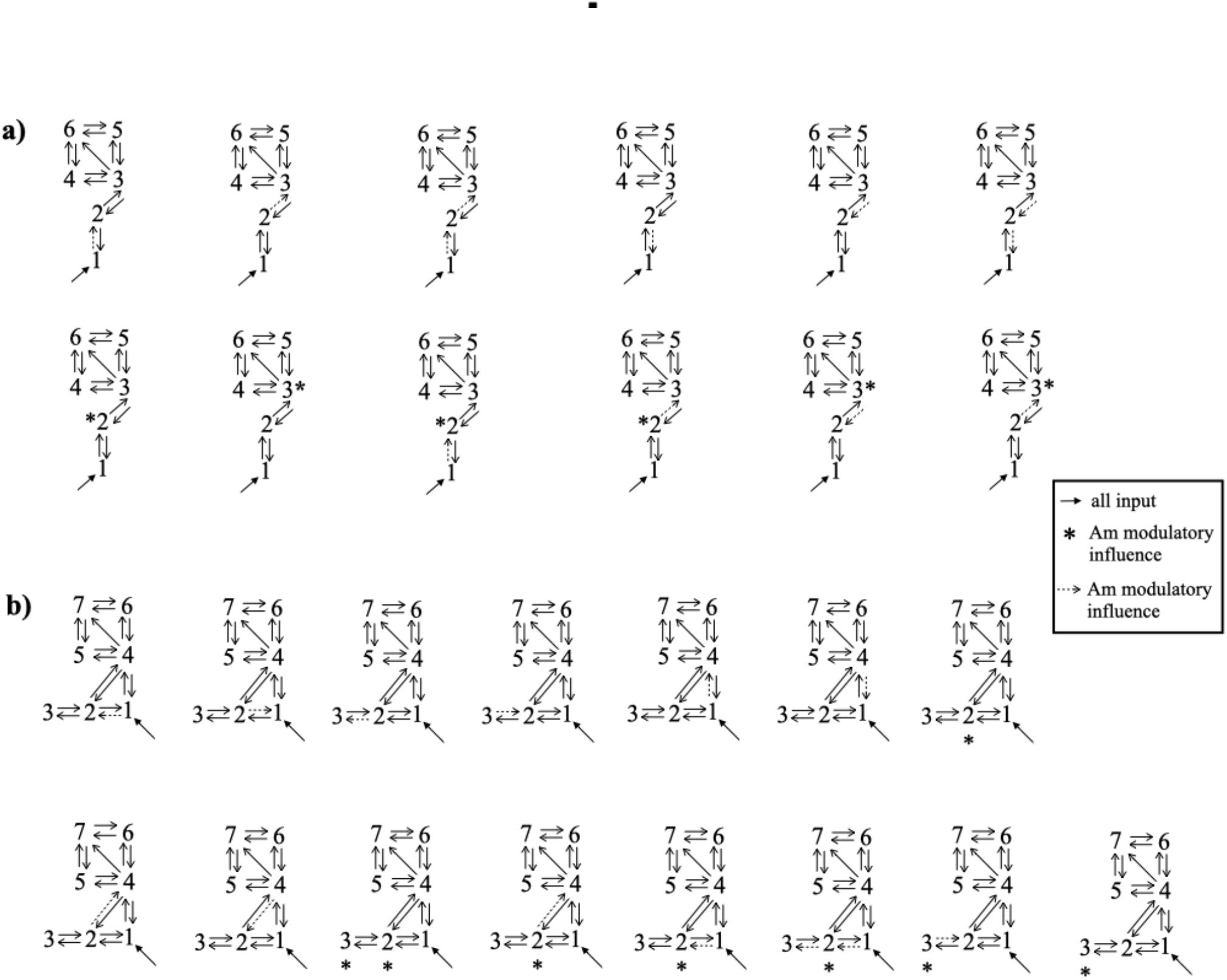
The three DCM families. Black dashed lines indicate input into the system by all conditions; black lines indicate fixed connectivity.

#### DCM Model comparison

Firstly, inference on model structure was performed using Fixed Effects Bayesian Model Selection (FFX BMS) (Penny et al., 2004), in order to compare the individual models over the BOLD sequence of interest. The model’s Free Energy, F, a lower bound of the model’s log-evidence, accounting for model complexity as well as data fit, was used to compare the likelihood of the different models to explain the data. Relative log-evidences, or differences in F, were converted into model posterior probabilities, p, indicating that the respective model/family has a probability p of being the best model explaining the data amongst all considered. Evidence was “strong” if p>0.95, which stands for a difference in F greater than 3, and “positive” if 0.75<p<0.95, which stands for a difference of F between 1 and 3 (Penny et al., 2004 and Penny et al., 2010).

Secondly, inference on the optimal model parameters was performed. The structure of the connectivity model was assumed to be the same for both sequences and a FFX analysis of the model parameter estimates was performed using Bayesian Parameters Averaging (BPA) (Acs and Greenlee, 2008, Garrido et al., 2007; Neumann and Lohmann, 2003).

## Results

### Self-report questionnaires

Clinical and demographical data are reported in table 1. No significant group differences were observed for STAI-T and BDI-II scores.

### Behavioral results

A repeated-measure ANOVAs of RTs to the key press and release, using the group (FND vs controls), prime type (aversive vs neutral), and conscious level (masked vs unmasked) as factors, revealed a main effect of group both for the key press [F=64.363 p < 0.00001] and the release [F=13.326 p < 0.00001] actions. The healthy controls were significantly faster than FND participants in both conditions (Fig. 3). No other main effects or interaction reached a significant threshold. An analysis of missing responses (i.e., slower than cutoff) also revealed a distinctive pattern in both groups. Both groups missed the release action more often than the key press action (i.e., in the requested temporal window), but in the FND group the number of missed responses was three times bigger than the key press misses (Tab. 2). Noteworthy, in FND participants, missed responses in the key press conditions were significantly associated with the aversive masked (p < 0.005) and aversive unmasked primes (p<0.014) in the key press condition (Tab. 2).

**Fig. 3:**
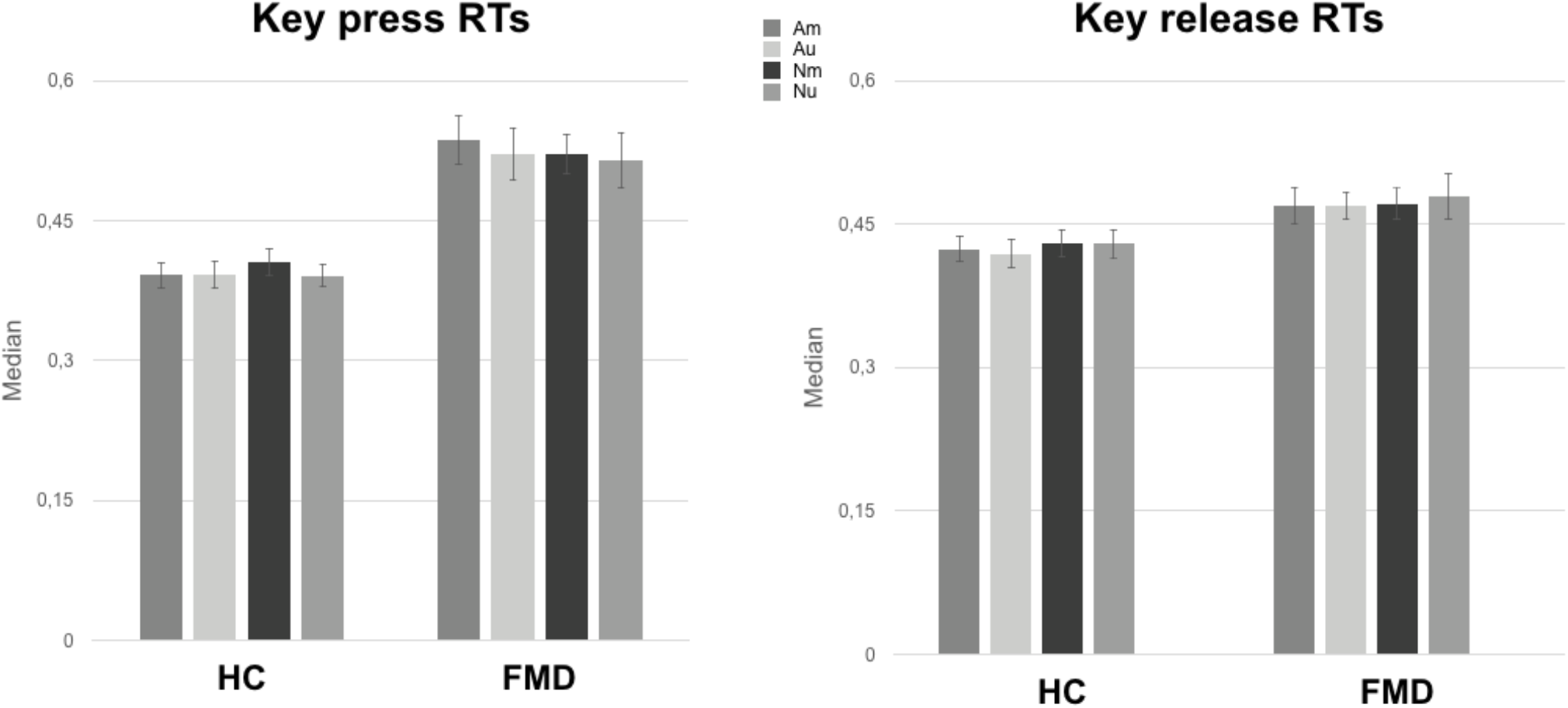
Median of reaction times. Error bars show confidence intervals level at the 95%.

**Table 2:** Missing responses.

|  | **FMD group** | **Healthy controls** | **Pearson Chi-Square** |
| --- | --- | --- | --- |
| **Am press** | 32 | 14 | 0.005 |
| **Au press** | 36 | 19 | 0.014 |
| **Nm press** | 35 | 23 | 0.085 |
| **Nu press** | 23 | 16 | 0.22 |
| **Am release** | 80 | 27 | 0.000 |
| **Au release** | 75 | 28 | 0.000 |
| **Nm release** | 77 | 26 | 0.000 |
| **Nu release** | 73 | 28 | 0.000 |

### Neuroimaging Results

#### Whole-brain analyses: Main effects

First, I tested for the main effect of the motor task, by comparing activations to the onset of all four primes [Au, Nu, Am, Nm] which summoned for action preparation, to the offset time of the landscape picture which corresponded to the interruption of motor action, for both groups together. As expected, action preparation produced higher activation in several premotor / planning areas (left SMA, right superior parietal lobule/SPL, left inferior frontal gyri/IFG/BA45/ BA47) and motor areas (bilateral precentral gyri, caudate, thalami, cerebellar lobules V and VIIa), as well as in the right periacqueductal grey (PAG), right precuneus, left posterior cingulate cortex (PCC), and fusiform cortex (all P < 0.05 FWE corrected) (Tab. 3). On the other hand, the release action produced greater activation in inhibitory control and attention-related areas including the right precentral gyrus, right angular gyrus, ACC, as well as the left precuneus and bilateral occipital cortex (P < 0.05 FWE corrected) (Tab. 3). The main effect of prime awareness was tested by comparing unmasked and masked conditions at the onset of the primes, again across both groups together. The unmasked (consciously seen) primes [Au+Nu>Am+Nm] activated significantly more the fusiform cortex, as well as bilateral parietal areas, bilateral IFG (pars opercularis), and left superior medial prefrontal cortex/mPFC (P < 0.05 FWE corrected) (Tab. 4). Conversely, the masked (unseen) primes [Am+Nm>Au+Nu] produced no significant effect above threshold. Finally, the main effect of emotional (aversive) primes [Am+Au>Nm+Nu] across both groups activated the right anterior insula, right amygdala, bilateral basal ganglia (including caudate and pallidum), plus bilateral mPFC, thalamus, and the cerebellar lobule VIIa (P < 0.05 FWE corrected) (Tab. 4). These activations indicate that aversive stimuli enhanced brain response not only in limbic areas, as expected, but also in several subcortical motor areas. Taken together, these results confirm that this task successfully recruited neural circuits associated with motor control and emotional reactions.

**Table 3:** Main effect of the task, P<0.05 FEW.

|  | MNI coordinates |  |  |  |  |
| --- | --- | --- | --- | --- | --- |
| Region | x | y | z | Ke | Z Scores |
| Primes > Key release |  |  |  |  |  |
| Left fusiform | -32 | -52 | -16 | 47813 | Inf |
| Right precentral gyrus BA9 | 40 | 6 | 32 |  | Inf |
| Left SMA BA6 | -2 | 6 | 52 |  | Inf |
| Left precentral gyrus BA6 | -46 | 0 | 38 |  | Inf |
| Left inferior frontal gyrus BA9 | -36 | 8 | 30 |  | Inf |
| Right superior Parietal Lobule BA7 | 30 | -52 | 44 |  | Inf |
| Left inferior frontal gyrus BA45 | -32 | 24 | 6 |  | 8.50 |
| Right caudate head | 4 | 10 | 4 |  | 8.10 |
| Left caudate | 0 | 16 | 4 |  | 8.07 |
| Left thalamus | -8 | -20 | 8 |  | 9.45 |
| Right thalamus | 8 | -16 | 8 |  | 9.04 |
| Right periaqueductal grey | 2 | -28 | -2 |  | 9.85 |
| Right precuneus | 16 | -64 | 40 | 48 | 9.41 |
| Left posterior cingulate cortex BA31 | -14 | -24 | 40 | 111 | 6.55 |
| Left insula BA13 | -40 | -4 | 14 | 43 | 6.41 |
| Left fusiform BA20 | -40 | -8 | -22 | 24 | 6.24 |
| Right middle orbitofrontal cortex BA11 | 28 | 36 | -12 | 143 | 6.00 |
| Right lobule VIIa crus I | 30 | -86 | -29 | 1599 | 7.49 |
| Left lobule V | -2 | -56 | -5 | 193 | 5.78 |
| Left lobule VIIa crus I | -40 | -72 | -29 | 410 | 5.15 |
| Key release > Primes stimuli |  |  |  |  |  |
| Left middle occipital gyrus | -18 | -98 | -2 | 945 | Inf |
| Right calcarine | 14 | -98 | -2 | 900 | Inf |
| Right superior temporal gyrus | 64 | -22 | 4 | 503 | Inf |
| Right precentral gyrus | 38 | -20 | 52 | 606 | Inf |
| Left superior prefrontal gyrus BA8 | -42 | 20 | 52 | 348 | 7.72 |
| Left middle temporal gyrus | -64 | -10 | -12 | 553 | 7.46 |
| Left medial temporal pole | -52 | 10 | -30 | 257 | 6.84 |
| Left angular gyrus | -44 | -70 | 34 | 86 | 5.55 |
| Left precuneus | -6 | -60 | 24 | 175 | 6.17 |
| Left ACC | -8 | 36 | -4 | 68 | 5.24 |
| Right angular gyrus | 50 | -64 | 32 | 46 | 5.22 |

**Table 4:** Main effect of the modality factor, P<0.05 FWE.

| MNI coordinates |  |  |  |  |  |
| --- | --- | --- | --- | --- | --- |
| Region | x | y | z | Ke | Z Scores |
| **Unmasked > Masked Prime across groups** |  |  |  |  |  |
| Left fusiform | -42 | -44 | -20 | 2209 | Inf |
| Right fusiform | 42 | -44 | -20 | 2154 | Inf |
| L IFG triangularis | -40 | 10 | 26 | 1202 | 6.49 |
| R IFG opercularis | 44 | 14 | 34 | 746 | 6.06 |
| Right IFG opercularis | 32 | 32 | -14 | 212 | 5.84 |
| Right angular gyrus | 34 | -54 | 42 | 178 | 6.21 |
| Left SPL | -30 | -56 | 40 | 118 | 5.73 |
| Left superior mPFC | -6 | 26 | 52 | 91 | 5.48 |
| **Aversive > Neutral Prime across groups** |  |  |  |  |  |
| Right anterior insula | 32 | 18 | -12 | 2089 | 5.65 |
| Left caudate head | -6 | 8 | -6 |  | 5.43 |
| Left caudate body | -6 | 6 | 12 |  | 4.19 |
| Right lateral global pallidus | 16 | 2 | -8 |  | 4.81 |
| Right amygdala | 20 | -2 | -12 |  | 4.44 |
| Left mPFC BA10 | -10 | 46 | 10 |  | 4.12 |
| Right mPFC BA10 | 4 | 48 | 12 |  | 4.80 |
| Right posterior mPFC | 10 | 20 | 56 | 1124 | 5.24 |
| Left IFG orbitalis | -34 | 22 | -10 | 177 | 5.31 |
| Right thalamus | 10 | -30 | -2 | 208 | 4.36 |
| Left VIIa crus I | -56 | -48 | -35 | 68 |  |

#### Whole-brain analyses: Group differences

Next, I tested for brain areas differently activated by the two groups as a function of the different prime conditions. To this aim, first an interaction analysis between groups and conscious awareness of primes was carried out. Masked primes elicited higher activation in the left PCC and cuneus, as well as in the left caudate, left thalamus, and anterior cerebellum in the healthy participants (P <0.05 FWE cluster-level corrected) (Tab. 5). On the other hand, the FND participants revealed a selective disengagement of the caudate, putamen, and vmPFC with the masked primes, an effect mainly driven by the aversive condition (Fig. 4 above). The opposite interaction (differential effects of unmasked primes) showed no significant activations. I also performed an interaction analysis between groups and the emotional value of primes (aversive vs neutral). The left ACC, right caudate, right putamen, bilateral vmPFC, and right cerebellar VIIIa in healthy participants were selectively more activated when exposed to aversive pictures (both the masked and the unmasked conditions) than to neutral pictures (P < 0.05 FWE cluster-level corrected). In contrast, these regions were relatively suppressed following aversive primes in the FND participants (Tab. 5, Fig. 4), particularly in the masked condition where both the vmPFC and caudate showed a marked reduction [Am trials] unlike other conditions and unlike controls. The opposite interaction showed no greater emotion effect in FND than HC. These whole-brain results therefore globally support this research hypothesis about a different emotion-action interaction between the two groups.

**Table 5:**
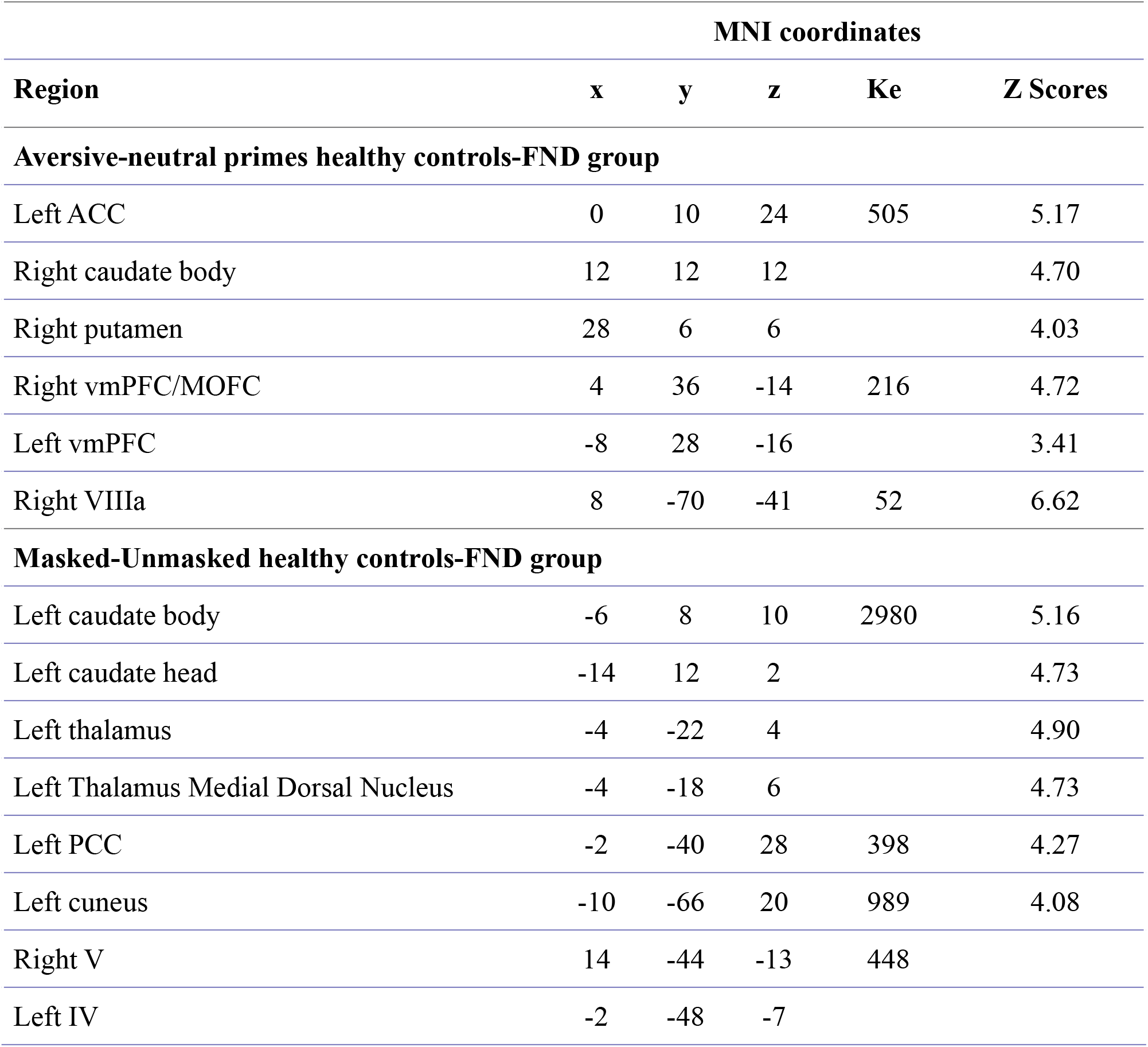
Interaction between type x groups and modality x groups, P<0.05 FWE cluster-leve.

**Fig. 4:**
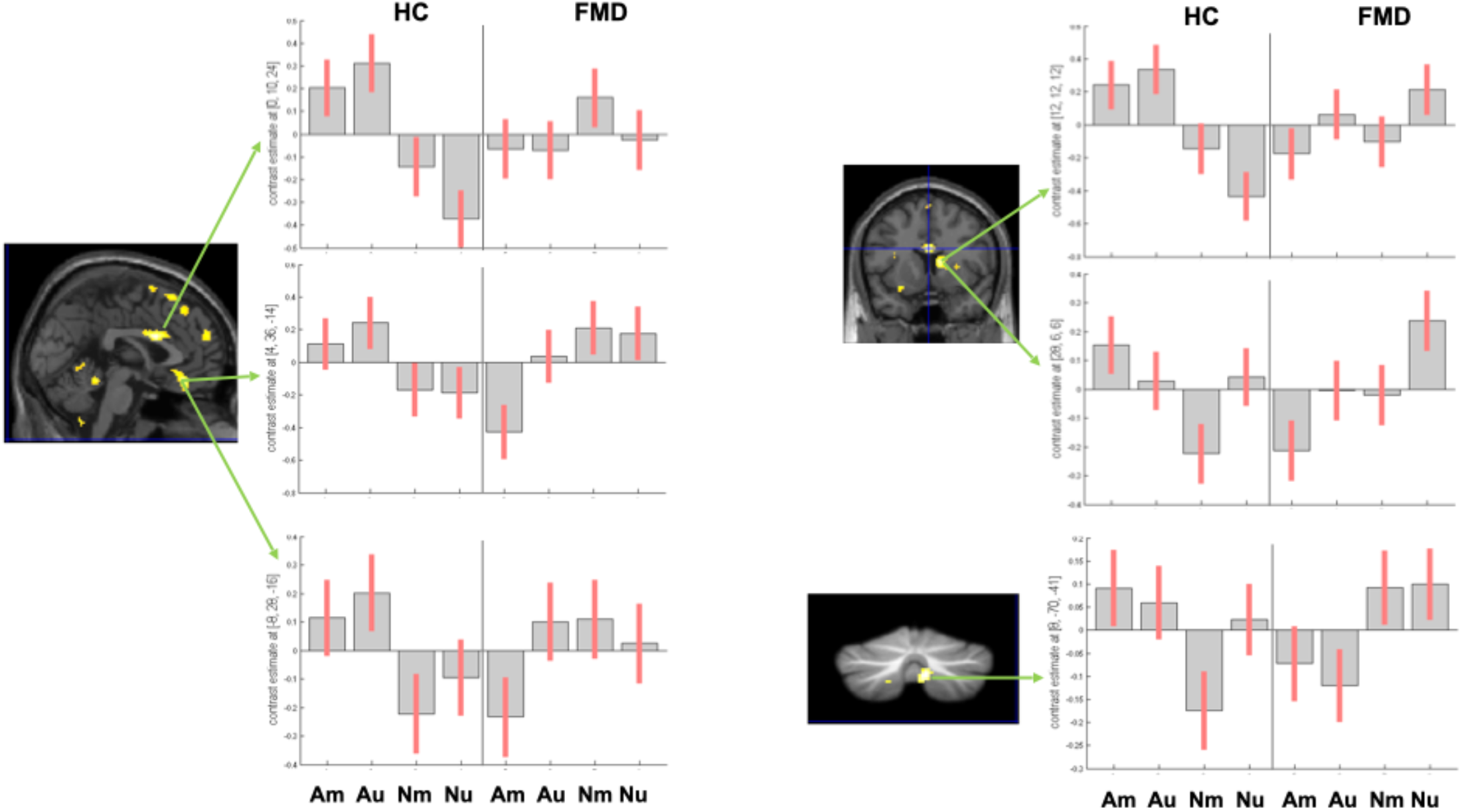
Differential brain responses in the two groups. Contrast maps and plots of beta estimates show fMRI results for the 2-way interaction analyses (group x aversive primes) in **a)** the left ACC (peak coordinates x= 0, y= 10, z= 24, P<0.05 FWE corrected) and the bilateral vmPFC (peak coordinates x= 4, y= 36, z= -14, x= -8, y= 28, z= -16, P<0.05 FWE corrected); **b)** the right caudate (peak coordinates x= 12, y= 12, z= 12, P<0.05 FWE corrected); the right putamen (peak coordinates x= 28, y= 6, z= 6, P<0.05 FWE corrected) and **c)** the right cerebellar lobule HVIIIa (peak coordinates x= 8, y= -70, z= 41, P<0.05 FWE corrected). (Mean ± 0.95 confidence intervals).

#### DCM Analyses

To address the main question of this study concerning functional interactions between brain areas, I applied DCM analyses to investigate the causal architecture of effective connectivity that may account for group differences during action control while being exposed to aversive masked and un-masked stimuli. DCM analysis were performed in each group separately to identify their distinctive functional network organization.

Among the ‘limbic’ family of networks, the “vmPFC” architecture was the winning model in the healthy control group when facing aversive masked prime (Fig. 5). The optimal model showed a positive modulation by the aversive masked condition on the left precentral gyrus, and a negative modulation on the backward connection from the left precentral gyrus to the left IFG. The log evidence was 138 with a posterior probability (Pp) of 1 in healthy controls, while in the FND group the log evidence was 0 with a Pp of 1.04e^-13^. This finding suggests a noteworthy pathway for information flow when healthy controls prepare action after being primed by an aversive scene at an automatic level. The DCM highlights a central input role for the vmPFC and an output node modulated by an aversive emotion in the left precentral gyrus, while the backward connections from left precentral gyrus to left IFG are directly inhibited by the masked aversive primes. The unmasked conditions produced no significant modulation of connections (Pp .38).

**Fig. 5:**
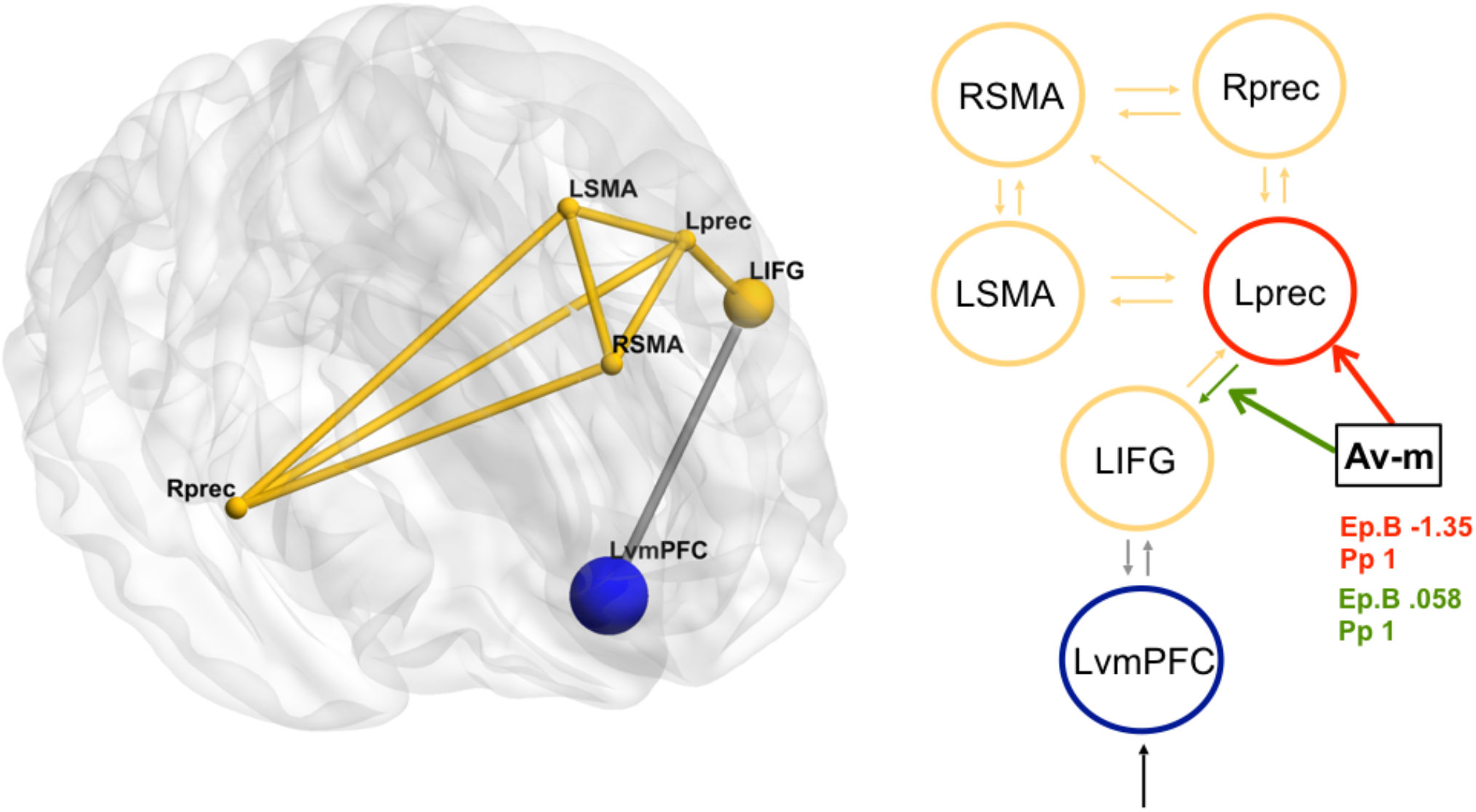
DCM winner models in healthy and FND groups. Black dashed lines indicate input into the system by all conditions; black lines indicate fixed connectivity; black bold lines and circles indicate the modulatory effect of masked aversive prime (Am). **Ep.B**: connection strength index; **Pp**: posterior probability.

On the other hand, in the FDM group, the winning model was the ‘self-focusing’ architecture, incorporating the “dmPFC-PCC” nodes instead of vmPFC (Fig. 6). The optimal model in this family also involved a reverse pattern of information flow. Now, the left IFG played the input role and its forward connection to the left precentral gyrus was positively modulated by the aversive masked condition (Log-e 90, Pp 1 in FND group vs Log-E 0 in the healthy group). The aversive unmasked condition produced no significant modulation. These findings suggest that in FND patients, the left IFG was the cortical region that drove information flow into the motor nodes.

**FIG 6:**
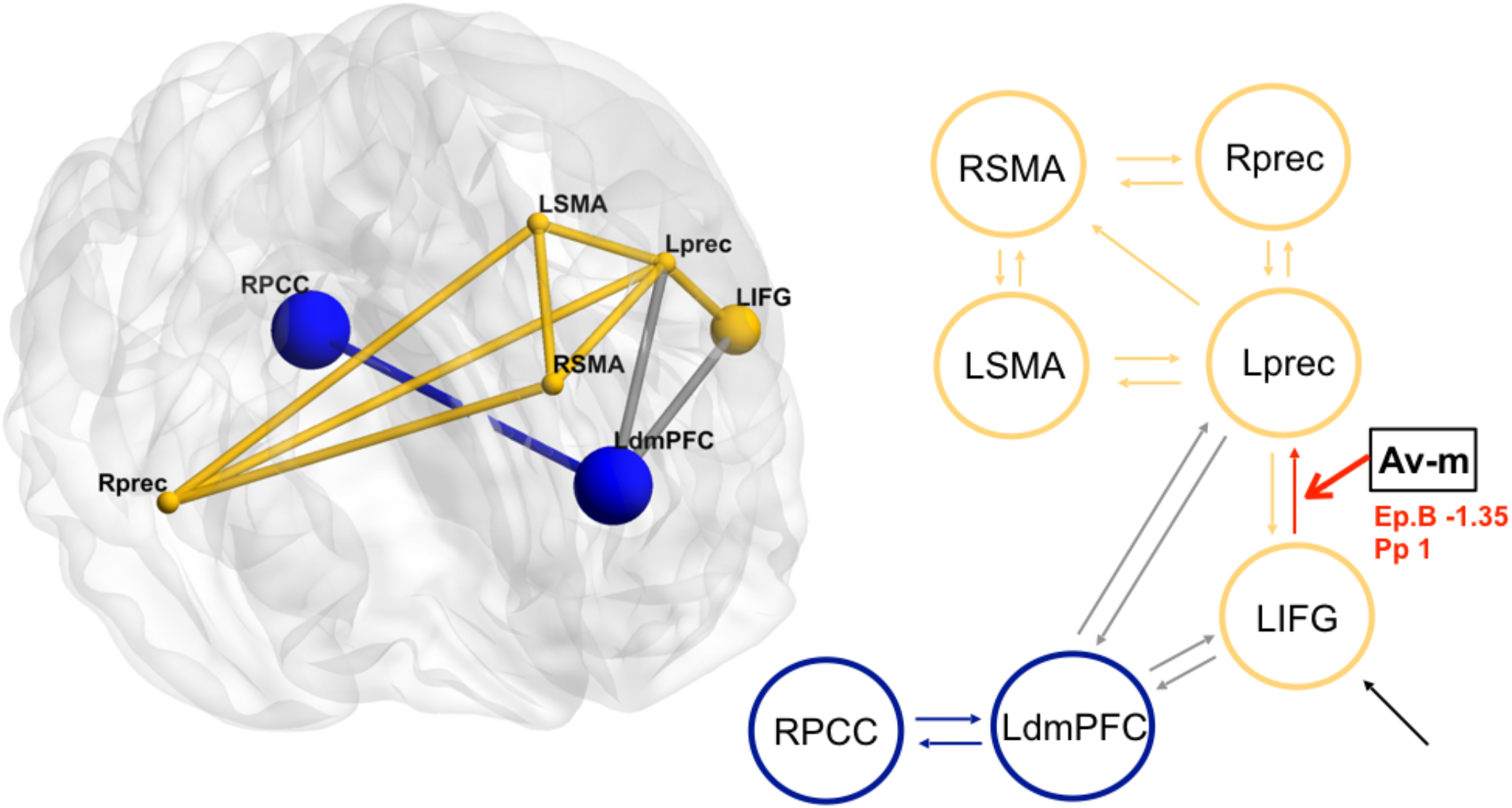

The third model family with a “vmPFC-amygdala” architecture had a very weak Pp across groups and conditions (Pp .38 in healthy group; Pp 0 in FND group).

Taken together, these results indicate that the two groups differed in the dynamic neural architecture of motor preparatory activity during our action priming task. Indeed, the “motor module” in our networks was differently hierarchically connected, with differential influences between the left IFG and the left precentral gyrus acting in an opposite direction between the two groups, selectively when exposed to masked (unseen) aversive primes.

Because of the significant number of missed key-releases in the FND group (i.e., in the requested temporal response window), we could not perform DCM analyses to compare the effective connectivity of these networks during the interruption of the movement (i.e. key release condition).

## Discussion

In the present study, I investigated whether FND patients and healthy volunteers would differ in their emotional reactivity to aversive primes, thus differently affecting their preparatory motor hand activity. The modeling framework here adopted makes the key assumption that the dynamics associated with a movement must partially depend on a rising urgency component that can be even independent of the perceptual ‘conscious’ (i.e. ‘seen’ in this context) evaluation process but rather being internally driven according to an automatic emotional reactivity. This study provides evidence of a peculiar neural architecture that links emotionality and hand preparedness in the FND participants from which the dynamics of the movement evolve to generate the resulting performance. Actually, at the behavioral level FND group performance differed significantly. They had a slow reaction time across all the task conditions and a high rate of missing key press responses associated to aversive primes.

The whole-brain analyses also supported the current hypothesis about a different emotion-action interaction between the two groups. Globally, motor preparation in response to the onset of visual stimuli (primes) activated a widespread network related to action planning and motor control (Valyear et al. 2015), at both the cortical (contralateral M1, SMA) and subcortical levels (basal ganglia and sensorimotor cerebellum consistent with the hand area), whereas motor interruption at the offset of the visual stimuli (key release) activated fronto-parietal networks related to attention and inhibition (ACC, IPL, IFG) as well as in the ipsilateral motor cortex. More critically, however, interaction analyses showed that the effect of emotional primes on motor preparation (both masked and unmasked) produced differential increases in the vmPFC, ACC, as well as subcortical motor relays (right caudate, putamen, and cerebellar VIIIa lobule) that were selectively inhibited in FND compared to healthy controls. These results are consistent with a central role of these regions in implementing and controlling motivated sensorimotor behaviors as well as with the urgency of commitment to a motor response (Thura and Cisek, 2017). Indeed, the vmPFC played its role in evaluating and controlling the perceived aversiveness in order to orient the behavioral response (Bzdok et al. 2013). These results support the a priori expectations of a greater involvement of vmPFC in healthy participants. On the other hand, the FND participants selectively did not engage the vmPFC and the related subcortical motor regions under the aversive conditions. It suggests a different motor commitment in the FND group. In addition, whole-brain analyses revealed also a different cerebellar involvement in the two groups. According to the role played by the cerebellum in the prediction of the sensory consequences of self-initiated actions (Cao et al. 2017), the healthy participants activated the anterior and sensorimotor cerebellum during the preparatory phase as a main effect of the group. Interestingly, the FND group engaged more posterior areas of the cerebellum, not engaged by motor planning activity but rather by emotional processing (Baumann et al. 2012).

More critically, the DCM analysis of brain dynamics during the task allowed to highlight distinct functional neural architecture linking emotionality to hand preparadness in the two groups of participants. A limbic family of networks was the winning in healthy controls, in which the motor/ premotor nodes received from vmPFC, and the left precentral gyrus (controlling right hand movements) constituted an output node directly modulated by the aversive nature of primes when masked. In turn, the left precentral gyrus appeared to have negative backward connectivity with the left IFG in the same prime condition. This result is consistent with an emotional modulatory effect on preparatory activity that may promote motor action. Thus, in the cortical dynamics controlling right hand movements, exposure to the masked aversive emotional information might drive faster recruitment of motor cortex to execute the anticipated action, e.g., pressing the key (Asfar et al. 2011; Churchland et al., 2010). Given also the better behavioral results performed by this group, these findings sustain that the emotional reactivity exerts a direct effect on the overt motor response, setting an initial condition of readiness to act.

In contrast, the FND group exhibited a different functional architecture of emotion-action interactions under the influence of emotional primes. In these patients, the winning family of networks was one connecting motor nodes with brain areas associated with self-referential focusing processes. Here, the optimal model revealed that the left IFG acted as the main driving input into the network and had an inhibitory forward connection to the left precentral gyrus that was enhanced by the masked aversive primes. Contrary to the control group, in the FND participants the aversive masked primes set a different dynamics in the hand action module, suggesting that another type of movement was then initialized. Given the association between FND group and missed responses in the aversive masked condition, I argue that the effect of the emotional reactivity on the overt motor response results in setting an initial condition from which the sequence of neural commands underlying were about to constrain the hand action and to do not act (Asfar et al. 2011; Churchland et al., 2010).

Taken together, the present findings unveil a twofold aspect about possible FNDs causation in terms of emotionality and voluntary motor control relationship. On one hand, FND patients may engage an abnormal self-focused processing mode when exposed to an aversive context that in turn interacted disruptively with the neural dynamic necessary to drive a willed movement. Evidently, the increased self-focused attention that misdirects the expected movement, as hypothesized by Edwards and colleagues, can be triggered by an emotional processing (Edwards et al. 2012). Moreover, such heightened self-focusing mode is consistent with the increasing evidence about an abnormal involvement of self-referential processing during voluntary emotional regulation in FND participants (Sojka et al. 2019). From a psychological point of view, this attitude is reminiscent of other psychological symptoms that also imply a heightened self-focused attention in their mechanism, as well as a sense of involuntariness and unwillingness. One example is the fear of blushing. Generally, self-focused attention does not only refer to a particular process (internally directed attention) but also to a particular kind of content about bodily states and/or one’s physical feature, such as, for example, to becoming red in the face or having awkward movements and sensations of body parts (Ingram, 1990; Bögels 1996). Accordingly, a bigger attention to physical precipitating factors (e.g., injury, illness) in FND has been reported by clinicians (Parées et al.2014b). Likewise, emotionality in FND patients may heighten their self-focused attention on proprioception and movement control under circumstances perceived as alarming (Arciero and Bondolfi, 2009).

On the other hand, it is remarkable that the FND group showed an abnormal pattern of emotion-action interaction in the presence of aversive scenes only when exposed in a subliminal fashion, namely in the masked condition. This suggests that abnormal engagement of affective and self-related processes may predominantly occur at a more automatic, non-conscious, level without an overt “cognitive” modulation. Such effect driven by an automatic process would be consistent with the subjective experience of FND patients who report a sense of involuntariness and unwillingness in the manifestation of their symptoms.

The current data allow us to add some knowledge to better understand the potential underlying FND process. First, they unveil the key role played by the emotional reactivity to initialize the dynamical system whose evolution will produce a movement activity. It highlights the strong and peculiar link between emotion and action in FN disorders. Second, the findings indicate that this processing occurs at a more automatic level. It makes clearer why FND patients report a sense of involuntariness and unwillingness of the manifestation of their symptoms. Third, I gather that the FND underlying emotional disorder can be referred to an abnormal self-referential focusing attitude. This attitude is associated with an external polarization trait that characterizes the personality of individuals affected by FND (Jaspers, 1964; Arciero and Bondolfi, 2009).

Due to the variability of FND manifestations, the main goal was to investigate the link between emotionality and motor preparation in an automatic fashion as a tentative to shift to an emotion-action frame of reference to grasp this core aspect of FNDs. According to this model framework, I found that a self-referential affective engagement can drive the motor down-stream setting an initial condition of the dynamics of the preparatory movement not moving. Further investigations are needed to better relate these activation patterns to specific motor symptoms, clinical history, and other psychological characteristics of individual patients. Nevertheless, these findings contribute to reduce the explanatory gap that exists regarding how emotionality might be translated into a potential process of FN symptoms production, showing how the FND participants’ affective engagement can be closely related to motor control brain activity and broadening the scope of the clinical intervention.

---

**Table S1:**
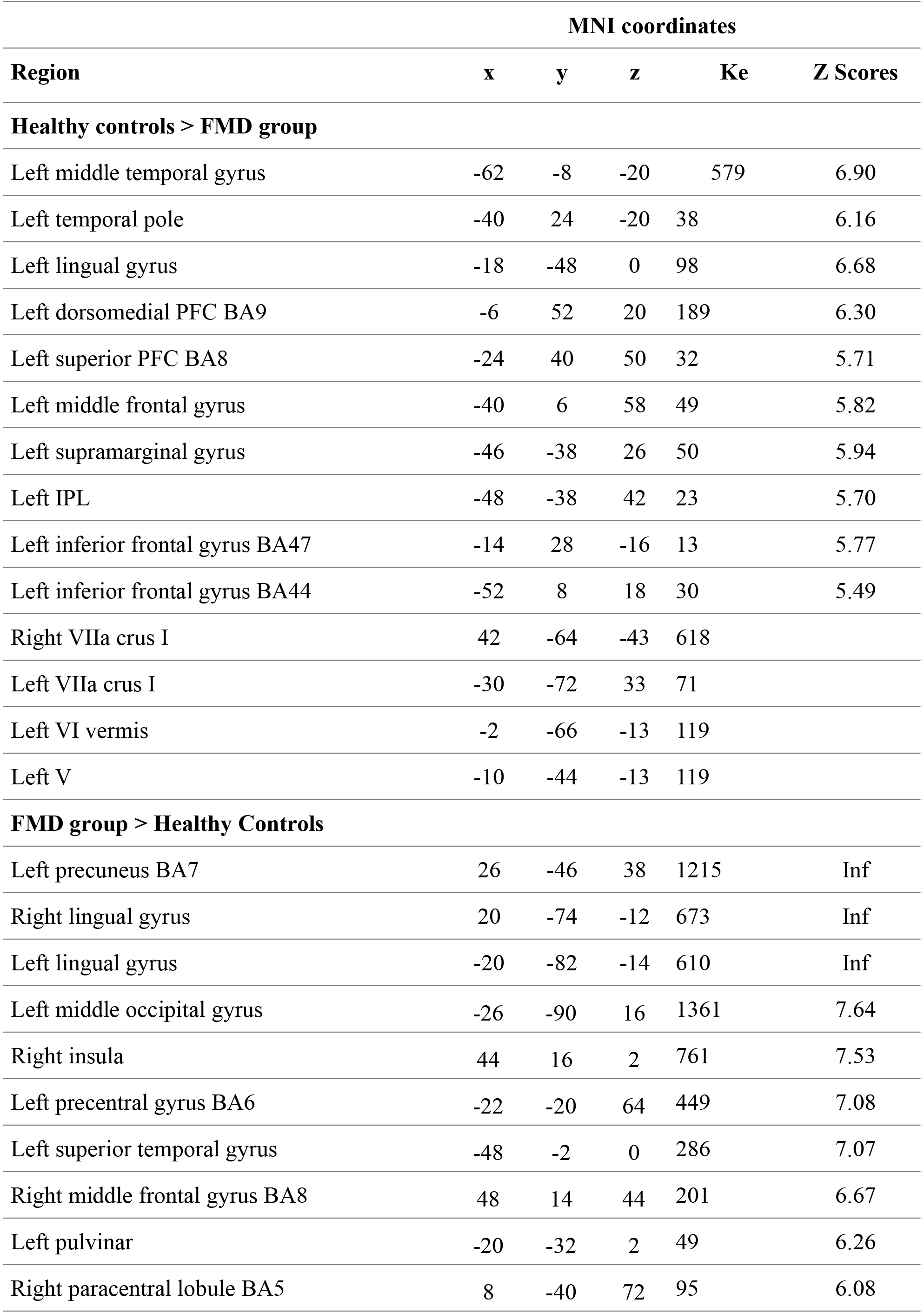
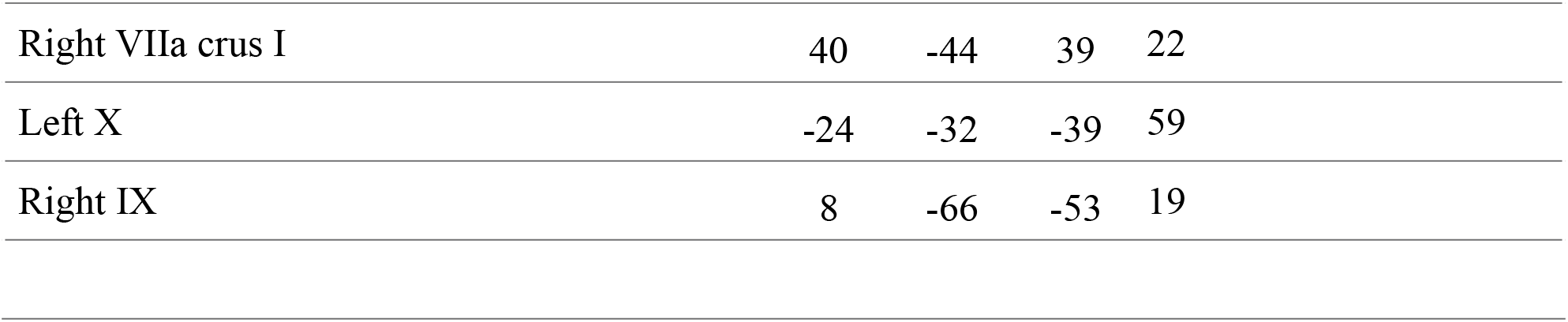
Main effect of the group, P<0.05 FWE.

|  | MNI coordinates |  |  |  |  |
| --- | --- | --- | --- | --- | --- |
| Region | x | y | z | Ke | Z Scores |
| Healthy controls > FMD group |  |  |  |  |  |
| Left middle temporal gyrus | -62 | -8 | -20 | 579 | 6.90 |
| Left temporal pole | -40 | 24 | -20 | 38 | 6.16 |
| Left lingual gyrus | -18 | -48 | 0 | 98 | 6.68 |
| Left dorsomedial PFC BA9 | -6 | 52 | 20 | 189 | 6.30 |
| Left superior PFC BA8 | -24 | 40 | 50 | 32 | 5.71 |
| Left middle frontal gyrus | -40 | 6 | 58 | 49 | 5.82 |
| Left supramarginal gyrus | -46 | -38 | 26 | 50 | 5.94 |
| Left IPL | -48 | -38 | 42 | 23 | 5.70 |
| Left inferior frontal gyrus BA47 | -14 | 28 | -16 | 13 | 5.77 |
| Left inferior frontal gyrus BA44 | -52 | 8 | 18 | 30 | 5.49 |
| Right VIIa crus I | 42 | -64 | -43 | 618 |  |
| Left VIIa crus I | -30 | -72 | 33 | 71 |  |
| Left VI vermis | -2 | -66 | -13 | 119 |  |
| Left V | -10 | -44 | -13 | 119 |  |
| FMD group > Healthy Controls |  |  |  |  |  |
| Left precuneus BA7 | 26 | -46 | 38 | 1215 | Inf |
| Right lingual gyrus | 20 | -74 | -12 | 673 | Inf |
| Left lingual gyrus | -20 | -82 | -14 | 610 | Inf |
| Left middle occipital gyrus | -26 | -90 | 16 | 1361 | 7.64 |
| Right insula | 44 | 16 | 2 | 761 | 7.53 |
| Left precentral gyrus BA6 | -22 | -20 | 64 | 449 | 7.08 |
| Left superior temporal gyrus | -48 | -2 | 0 | 286 | 7.07 |
| Right middle frontal gyrus BA8 | 48 | 14 | 44 | 201 | 6.67 |
| Left pulvinar | -20 | -32 | 2 | 49 | 6.26 |
| Right paracentral lobule BA5 | 8 | -40 | 72 | 95 | 6.08 |

|  |  |  |  |  |
| --- | --- | --- | --- | --- |
| Right VIIa crus I | 40 | -44 | 39 | 22 |
| Left X | -24 | -32 | -39 | 59 |
| Right IX | 8 | -66 | -53 | 19 |

